# Rapid ecosystem collapse and biofilter formation following seabed methane leakage

**DOI:** 10.64898/2025.12.05.692476

**Authors:** Qianyong Liang, Longhui Deng, Xinyue Liu, Xi Xiao, Ruize Xie, Jialin Hou, Jing Wang, Weikang Sui, Ningyuan Lu, Zian Tong, Danyue Huang, Yanwei Wang, Yingchun Han, Jing Zhao, Binbin Guo, Wei Zhang, Minghui Geng, Tianxin Ren, Wenqi Ye, Zheng Xiong, Liang Dong, S. Emil Ruff, Christof Meile, Jun Tao, Xiyang Dong, Fengping Wang

**Author notes:** These authors contributed equally to this work (Qianyong Liang, Longhui Deng). Correspondence: Fengping Wang, Xiyang Dong, or Longhui Deng.

## Abstract

Globally, vast amounts of methane are trapped within subseafloor gas hydrates. Recent evidence suggests that climate- and human-induced perturbations can destabilize gas hydrates and trigger methane release, yet the response of deep-sea ecosystems to these changes remains poorly understood. Here we show multi-year *in situ* monitoring of seabed ecosystem following methane leakage induced by hydrate exploration activities. Within just two years, benthic microbial and eukaryotic diversity in the affected areas declined significantly, while microbial and macrofaunal abundance increased. Integrated geochemical and omics analyses reveal the rapid successional shift of seabed ecosystem to a novel chemosynthetic system. Aerobic methanotrophs (*Methyloprofundus*) co-established with unexpectedly fast growing anaerobic methanotrophic archaea (ANME-3), accompanied by a rapid recruitment of opportunistic polychaetes that bioirrigated the seabed to >50 cm depths. The active animal-microbe interactions sustained exceptionally high rates of methane oxidation that utilized diverse electron acceptors. We demonstrate that methane hydrate destabilization can trigger rapid collapse of native seabed ecosystem while driving the formation of an effective ‘methane biofilter’ consuming this potent greenhouse gas much faster than previously estimated. Understanding seabed ecosystem response and feedback is critical for predicting benthic methane cycling under ongoing global change and for developing sustainable strategies for methane hydrate resource management.

## Main

Marine sediments hold Earth’s largest reservoir of methane, a clean fossil energy resource and potent greenhouse gas^1^. Methane is stored in sediment in its dissolved and gaseous forms, or as solid gas hydrate under low temperature (<4℃) and high pressure (>60 bar), a crystalline precipitate of water and hydrocarbons that harbors 500-2,000 gigatons of carbon (Gt C) worldwide^1^. Mounting evidence indicates that methane efflux from seabed is greater than previously recognized^2–5^, a trend likely to be exacerbated by global change^6–8^. Climate-driven increases in bottom-water temperature^7,9,10^, coupled with expanding oceanic resource exploration and mining activities^11,12^, threaten to destabilize hydrate reservoirs and enhance the advective flux of methane. This may lead to increasing, yet unconstrained amounts of methane escape to the upper hydrosphere or even atmosphere^13–15^. Hydrate dissociation-driven emission of methane was estimated to add 0.4–0.5°C to the warming initially caused by anthropogenic activities over millennial timescales^16^. Aerobic oxidation of methane released from seabed consumes oxygen in the seawater, together with the warming-induced reduction in oxygen solubility and ocean ventilation, might lead to expanding ocean hypoxia^17,18^.

Marine benthic ecosystems form a critical biological barrier — the “methane biofilter” — that effectively mitigates methane emission from the seabed^19,20^. The efficiency of this filter hinges on complex microbial and faunal communities sustained by methane seepage^21,22^. Along global continental slopes, the deeply sourced methanerich fluids discharge at seafloors, forming ‘cold seeps’ that sustain some of the most prolific chemosynthetic ecosystems on Earth^19^. Cold seep biota consume up to 100% of the upward transported dissolved methane locally, although only covering less than 0.05% of the slope area, they account for 10-20% of the total methane oxidation at continental slopes, representing a globally relevant gatekeeper to restrict seabed methane emission^22,23^. A fundamental knowledge gap, however, is the timescale of biofilter development following a sudden methane influx. The anaerobic methanotrophic archaea (ANME) central to effective benthic methane removal exhibit cell doubling times of months to years^24–29^, while the successional development of the requisite macrofaunal communities, which mediate methane consumption through their interactions with microorganisms, occurs over decades to centuries^19,30–37^. Thus, while direct monitoring of the *in situ* response of seafloor biota to abrupt methane influx remained lacking, the time span of forming an efficient methane biofilter following methane leakage has been considered rather long and proposed as a “time window”, during which large quantities of methane can be liberated from the seafloor to upper ocean or even atmosphere^14,25^.

Here, we report a six-year *in situ* study (2018–2023) that directly tracks the response of seabed ecosystem to an abrupt methane leakage event triggered by gas hydrate exploration in the South China Sea (SCS, Fig. 1a). We tracked the spatiotemporal changes of seabed communities and sediment geochemistry in the leakage-impacted areas annually (Fig. S1), using advanced underwater cameras, acoustic instruments, chemical sensors, and sampling facilities, combined with omics-based analyses, geochemical modeling, and isotopic rate measurements in the laboratories. We further compared the newly formed chemosynthetic ecosystem (hereafter called “Newborn Seep”) with both adjacent and distant non-seep and mature seep sites in the SCS (Fig. S2). Contrary to the established view of a slow biological response, our results demonstrate the prompt emergence of a distinct chemosynthetic ecosystem capable of efficient methane filtration at the initial response stage.

**Fig. 1.**
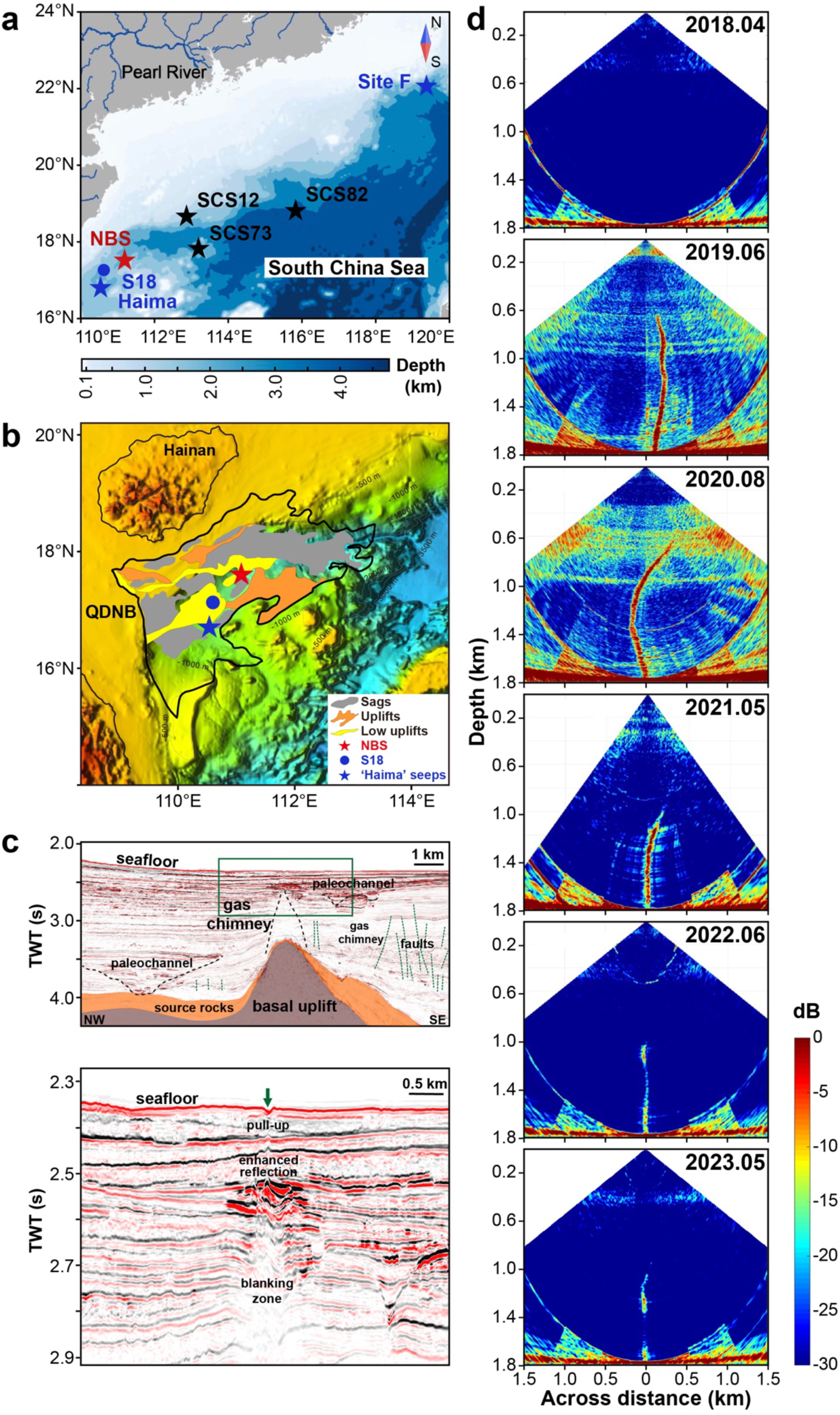
Tracking methane leakage triggered by gas hydrate drilling. **(a)** Map showing bathymetric features and sampling sites in the South China Sea (SCS), including the “Newborn Seep (NBS)”, mature cold seeps (S18, ‘Haima’ and ‘Site F’ cold seeps), and non-seep continental slope sites (SCS12, SCS73, SCS82). Note that two non-seep background sites 19-NS-BG1 and 19-NS-BG2 were taken ∼200 m away from the discharge center of NBS in 2019, before the spreading of leakage impact on seafloor (Fig. S1). **(b)** Geological setting of the study region within Qiongdongnan Basin (QDNB) of the SCS. **(c)** Seismic reflection profiles showing shallow subsurface structures, fluid pathways, and enhanced reflections. TWT: Two-way traveltime. **(d)** Water-column acoustic images from April 2018 to May 2023, revealing recurring acoustic flares (high-backscatter zones) indicative of ongoing gas bubble release from the hydrate-bearing seabed.

## Methane leakage triggered by gas hydrate drilling

The study site is located in Qiongdongnan Basin of the SCS at a water depth of 1,766 m, where gas hydrate inventory was estimated to reach ∼ 6.5×10^9^ tons of carbon over an area of ∼6×10^4^ km^2^ (Fig. 1a, b)^38^. High-resolution 3D seismic data reveal that this site is connected to a subseafloor gas chimney that originates from a basal uplift, appearing as a seismic blanking zone with base width of ∼1 km and narrows upwards to less than 0.2 km (Fig. 1c). Above the gas chimney, the pull-up enhanced reflections indicate the presence of highly saturated gas hydrates within the leakage pathway. In 2018, a drilling expedition was conducted to assess the gas hydrate resource^39^. Before drilling, the seafloor at the drill site appeared as ‘bare’ sedimentary floor with scarce benthic fauna and no gas plume (Fig. S2). A borehole in diameter of ∼22 cm was drilled using the logging tools from Schlumberger and subsequently sealed with heavy active mud^39^. While no gas discharge was observed after sealing the borehole in 2018, a distinct gas plume was detected by multibeam echosounders since 2019, suggesting the initiation of methane leakage during 2018–2019 (Fig. 1d). The plume reached ∼1.2 km and ∼0.8 km above the seafloor during 2019–2020 and 2021–2023, respectively. Elevated methane concentrations (up to 10 μmol L^-1^) were detected by underwater sensors in water column above the discharge center, indicating that those gas bubbles were rich in methane (Fig. S3).

## Rapid ecosystem collapse and restructuring

The seabed ecosystem at the leakage site underwent a rapid and dramatic transformation. Within two years of the leakage onset, bacterial and archaeal richness in surface sediments (0–100 cm) within 20 m of the discharge center declined by >50% (Fig. 2a). A concurrent decline in eukaryotic richness and all other diversity indices (Chao1, Shannon, and Simpson) marked a systematic ecological collapse of the native seabed ecosystem (Fig. S4). Following the initial decline, microbial richness appeared to rebound since 2020, indicating a rapid seabed ecosystem restructuring. In contrast, microbial abundance remained stable or even increased following the methane leakage, suggesting an immediate, physiological response of seabed microbial community to abrupt methane input. Concurrently, the seafloor morphology was radically altered. The previously barren sediment transformed into an extensive ‘worm bed’ within one year from 2020 to 2021 (Fig. 2b), associated with the rapid increase of opportunistic fauna dominated by Spionid polychaetes, Acrocirrid polychaetes, and Metridinid copepods (Fig. S5A). Bayesian δ¹³C mixing model calculations further show the substantial contributions of methane-derived carbon (20-70%) to the faunal diets (Fig. S5B). The observed suspension- and/or deposit-feeding behaviors of these dominant fauna suggest that prey-predator interactions, instead of symbiosis that is prevalent in mature cold seeps, drove the highly efficient methane-based trophic transfer. Noteworthily, dense populations of opportunistic polychaetes observed in natural seeps worldwide had long been proposed as indicator of early-stage seep ecosystems^33–35,40,41^, a notion now directly corroborated by our post-leakage monitoring.

**Fig. 2.**
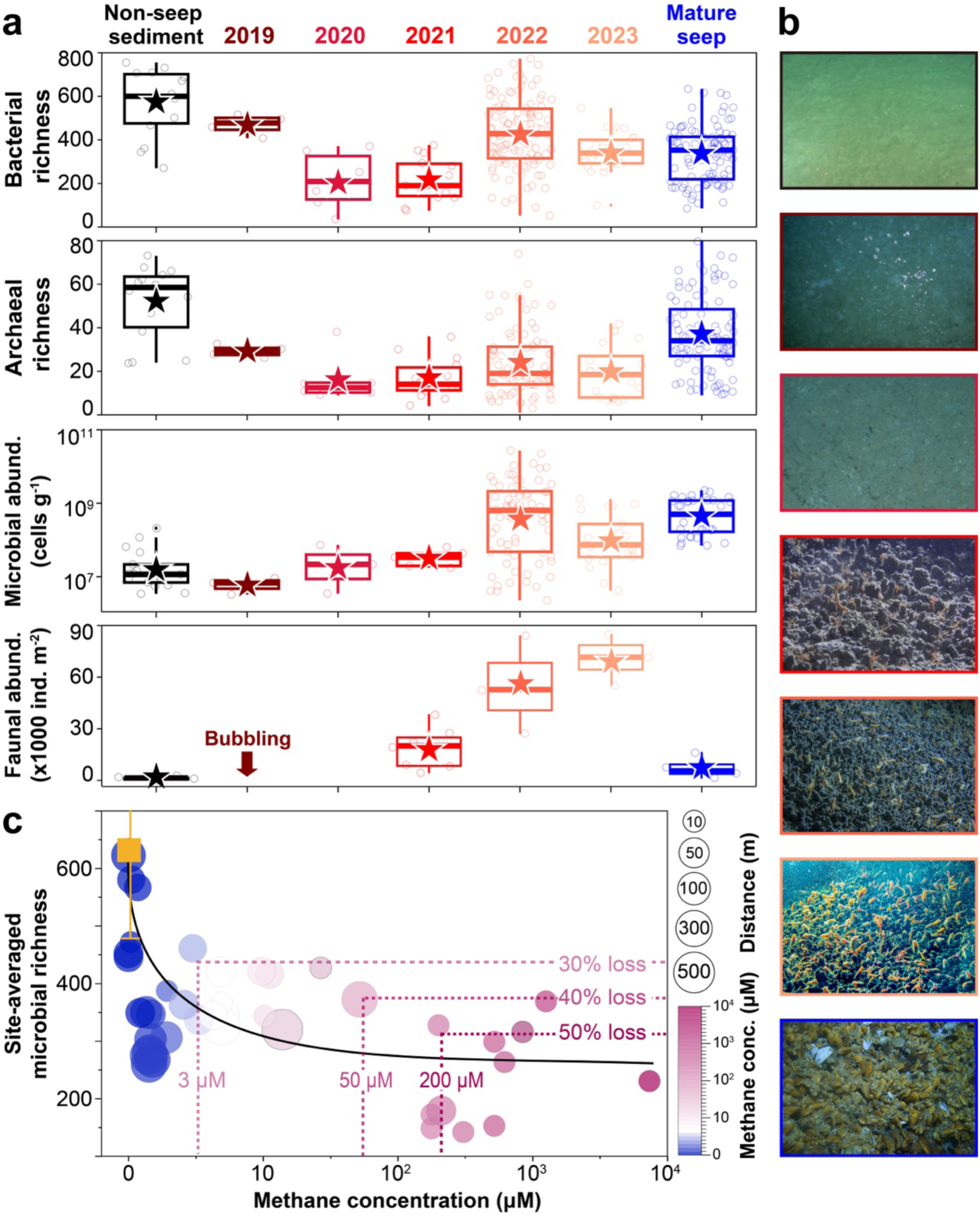
Ecological changes of seabed ecosystem following methane leakage. **(a)** Variations of bacterial and archaeal richness, microbial and macrofaunal abundances. Bacterial and archaeal richness were calculated as the quantities of amplicon sequence variants (ASVs) based on 16S rRNA gene sequencing. Microbial abundance was quantified based on qPCR of the bacterial and archaeal 16S rRNA gens. Macrofaunal abundance was determined by combining the sieving-based method and image analysis. The ‘non-seep sediment’ group includes only data from the adjacent non-seep background sites, while the ‘mature seep’ group includes data from the ‘Haima’ and ‘Site F’ cold seeps in the South China Sea. Boxplots show means (asterisks), medians, quartiles, and data ranges (same below). **(b)** Representative seafloor images of the pre- and post-drilling seafloors, as well as typical mature seep ecosystem (mussel bed) at ‘Haima’ cold seeps. **(c)** Negative correlation between the site-specific microbial richness and methane concentration. The site-specific values are calculated as the mean values across different sediment depths at each site. The symbol size and color indicate the distance to discharge center and methane concentration, respectively. The black solid line denotes the non-linear regression (adjusted R^2^: 0.4, *p*<0.001). The yellow symbol and error bar indicate the microbial richness in the adjacent non-seep background sites.

The impact of methane leakage expanded rapidly, affecting the wider seabed ecosystem within 2–3 years. Extensive sampling at 40 sites surrounding the discharge center revealed the occurrence of elevated methane concentrations (>10 folds of non-seep background values) in surface sediments even at 500-m distance from the discharge center (Fig. 2c). This was accompanied by a >30% reduction in microbial richness, demonstrating a remarkably rapid lateral propagation of the leakage impact (Fig. 2c). Synthesis of data from all sites reveals a significantly negative correlation between methane concentration and microbial richness (Fig. 2c, non-linear regression model F_1,39_= 25.23, *p*<0.001), matching our previous field observation^42^. Based on this correlation, methane concentrations of >3 μM corresponded to at least 30% microbial richness losses in surface sediments. While this correlation suggests that even micromolar increases of sediment methane concentrations can trigger significant microbial diversity changes, further research is needed to clarify whether these trends and thresholds are generally applicable to broader seabed ecosystems.

## Rapid assembly of a methanotrophic microbiome

Microbial community analysis revealed a rapid and non-stochastic succession towards a methanotroph-dominated ecosystem after methane seepage (Fig. 3). Pronounced shifts in dominant microbial taxa were observed even at class or phylum level: from bacterial *Alphaproteobacteria*, *Dehalococcoidia*, and *Anaerolineae*, and archaeal *Nitrososphaeria*, *Nanoarchaeia*, and *Bathyarchaeia* in the non-seep sediments, to bacterial *Gammaproteobacteria*, *Desulfobacteria*, and *Bacteroidia*, and archaeal *Methanosarcinia* in the Newborn Seep sediments (Fig. S6, S7). Microbial community succession rates, quantified as the temporal changes of microbial community distance (dissimilarity), were markedly high during 2019–2020 (Stage I: 0.2-0.3 yr^-1^, equivalent to a community turnover time of 3-5 years), but declined significantly during 2021-2023 (Stage II: <0.01 yr^-1^; Fig. 3a, b; Fig. S8). Matching this trend, the number of shared amplicon sequence variants (ASVs) between the non-seep and Newborn Seep communities decreased sharply during Stage I but rebounded slightly during Stage II. These trends indicate that the majority of the microbial succession occurred within the first 2-3 years after methane leakage. Although microbial community structures in the Newborn Seep sediments remained distinct from those in mature seeps (PERMANOVA, *p*<0.01), they rapidly progressed toward the mature seep community type (Fig. 3b). Community structures of some samples from Stage II already resembled those from mature seeps, indicating that the assembly of a typical seep microbial community can occur quickly within few years (Fig. 3b; Fig. S8, S9).

**Fig. 3.**
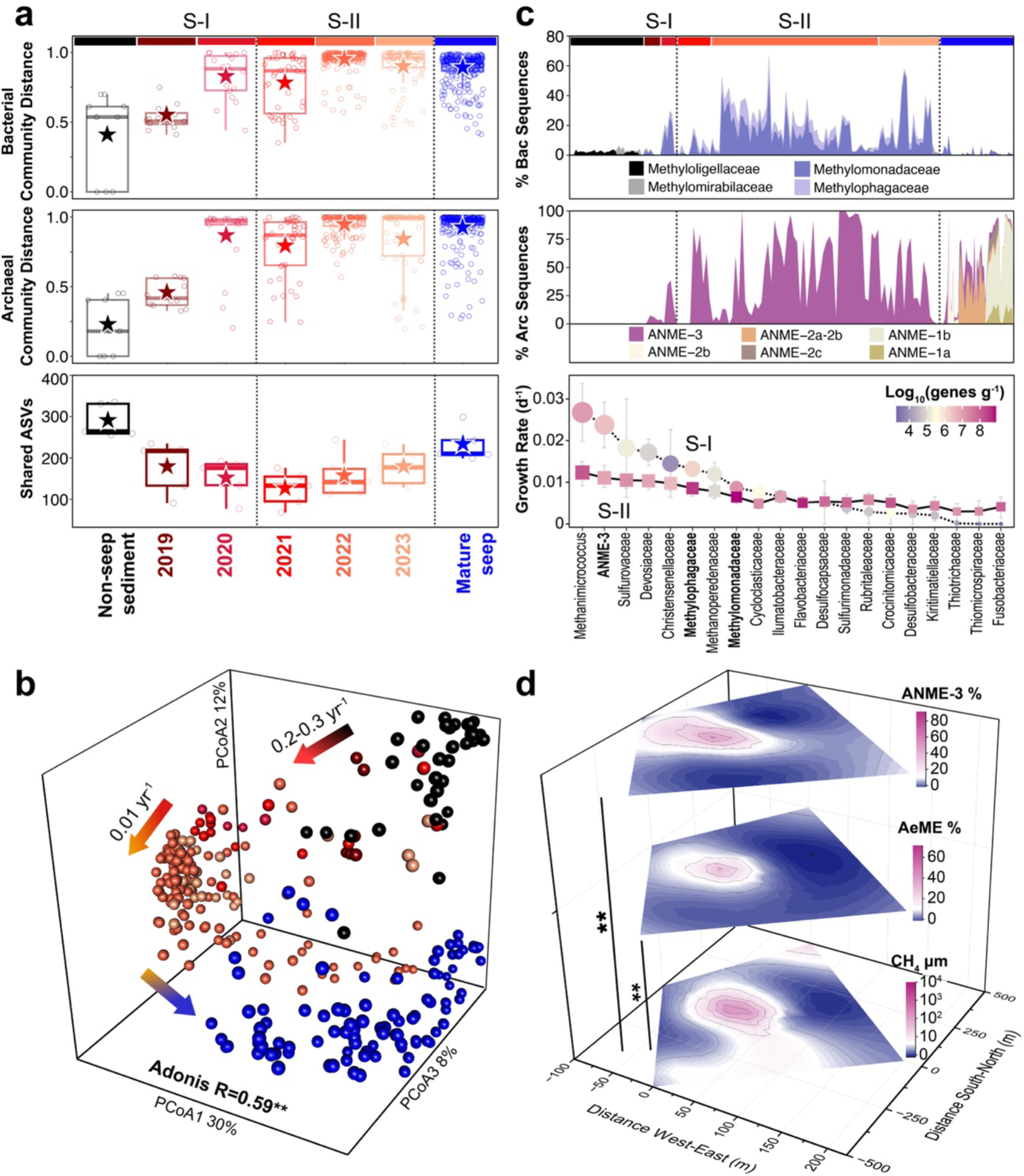
Spatiotemporal variations of sediment microbial communities in response to methane leakage. **(a)** Boxplots showing shifts in bacterial and archaeal Bray-Curtis community distance (dissimilarity) and shared amplicon sequence variants (ASVs) in the Newborn Seep and mature seep sediments, compared with the non-seep background sediments. Stage-I (S-I) and Stage-II (S-II) are defined based on the distinctly different community succession rates. **(b)** Principal coordinates analysis (PCoA) plots based on bacterial 16S rRNA gene sequences, showing the distinct community clusters across different seep development stages and years. Each point represents an individual sample, with the same color code shown in Fig. 3a. Adonis R and *p*-values indicate the proportion of variation explained by seep development stage (**: *p*<0.01). Similar trends for archaeal communities are shown in Fig. S8. **(c)** Variations in relative sequence abundances of dominant aerobic and anaerobic methanotrophic taxa across different seep development stages and years (upper and middle panel). The growth rates of different microbial lineages during Stage I and Stage II, respectively, were estimated as the log2-fold changes of 16S rRNA gene copies over time (lower panel). The symbol size is proportional to the growth rate, while symbol color indicates the average 16S rRNA gene abundances during Stage I or Stage II. **(d)** Integration of measurements on >40 sediment cores sampled around the discharge center from 20192023, revealing the spatial gradient of methane concentrations and relative abundances of ANME (mainly ANME-3) and aerobic methanotrophs (AeME, mainly *Methylomonadaceae* and *Methylophagaceae*). Methane concentrations were log_10_(x+1) transformed to better visualize the low-range values. Spearman correlations were performed to examine correlations of methane concentrations with the relative abundances of ANME-3 and AeME (**: *p*<0.01).

Remarkably, the presumably ‘fast-growing’ aerobic methanotrophs and ‘slow-growing’ anaerobic methanotrophs proliferated synchronously since 2019, and both became highly abundant in 2022-2023 (Fig. 3c). The changes of absolute 16S rRNA gene abundances over time suggest that dominant aerobic methanotrophs (*Methylomonadaceae,* up to 60 % of bacterial sequences) and anaerobic methanotrophs (ANME-3, up to 99 % of archaeal sequences) both reached quasi-stationary phases of growth in 2-3 years (Fig. S10). The estimated *in situ* doubling time of *Methylomonadaceae* (mainly *Methyloprofundus*) were 80-107 days (Fig. 3c), matching the previous estimates for aerobic methanotrophs in marine habitats^43^. Unexpectedly, ANME-3 exhibited a notably shorter doubling time (24-38 days) during Stage I, suggesting the even faster initial response of anaerobic methanotrophs than aerobic methanotrophs to methane leakage. This timescale is also shorter than the previously estimated doubling time of 100-200 days for ANME-3 in the newly erupted mud volcano sediments^43^, as well as the model-based prediction of 60-100 years for establishing a steady-state AOM community in response to new methane input^14,25,44^, challenging the paradigm of anaerobic methanotroph’s delayed response to sudden methane influx^14,24,25^. The unexpectedly rapid response of ANME-3 presents a compelling enigma. This archaeon is understood to canonically mediate sulfate-coupled AOM, a process with a low energy yield (∼ -25 kJ mol⁻¹ CH₄) that constrains growth rates^45^. Our observed doubling times challenge this thermodynamic expectation, raising a fundamental question: how does ANME-3 achieve such rapid proliferation? Substantial recruitment of ANME-3 from the surrounding sediments and waters can be excluded due to its extremely low abundance or absence in these environments (Fig. 3c). On the broader spatial scale, methane concentration gradients correlated positively with the relative sequence abundances of aerobic methanotrophs and ANME-3 (Fig. 3d, Spearman *p*<0.01), further supporting that elevated methane supply directly amplifies *in situ* methanotrophic responses.

## Effective methane removal by benthic biofilter

Time-series geochemical profiling within 20 m of the discharge center suggests a rapid methane removal from surface sediments in 2-3 years after methane leakage (Fig. 4). In 2019, elevated methane concentrations (10^2^-10^3^ μM) were detected within the 0-100 cm sediments, while isotopic signatures of dissolved inorganic carbon (δ^13^C-DIC), sulfate and ∑H_2_S concentrations remained at non-seep background levels, indicating the absence of active methane oxidation in 2019 (Fig. 4a). Yet starting from 2021, in parallel with the increasing faunal and microbial population density, methane oxidations were evidenced by a sharp concentration decrease to 1-10 μM and a negative shift of δ^13^C-DIC (-10‰ to -30‰) in the upper 40-50 cm sediments, where dense faunal burrows observed (Fig. 4a, b). In these strongly bioturbated layers, elevated total organic carbon (TOC) with highly negative δ^13^C-TOC co-occurred with net consumptions of oxygen, sulfate, and nitrate (Fig. 4b). Potential coupling of methane oxidation with nitrate reduction was also verified by isotopic labeling experiments, further suggesting the active methane oxidation using diverse electron acceptors. Whereas in the deeper non-bioturbated layers (50-100 cm), a pronounced δ^13^C-DIC depletion (-55‰ to -20‰) coincided with the sulfate reduction (16-20 mM) and ∑H_2_S accumulation (10^2^-10^4^ μM). Together these changes reveal effective methane consumptions throughout the top 100 cm of sediment at the Newborn Seep across two distinct geochemical zones: 1) active methane oxidations co-occurred with diverse redox reactions in the 0-50 cm sediment, likely promoted by the bioturbation-induced redox fluctuations, and 2) anaerobic methane oxidation coupled with sulfate reduction in the 50-100 cm sediments. The integrated methane oxidation fluxes throughout 0-100 cm sediments of the Newborn Seep, based on the time-dependent 1D reaction-transport model, were 30-40 mmol/m^2^/day, matching fluxes estimated using the Monod biomass-explicit model (30-60 mmol/m^2^/day). This budget accounts for 60-100% of the steadystate methane oxidation flux under mature seep settings (∼50 mmol/m^2^/day), further supporting the rapid establishment of effective methane biofilter within a few years after methane leakage.

**Fig. 4.**
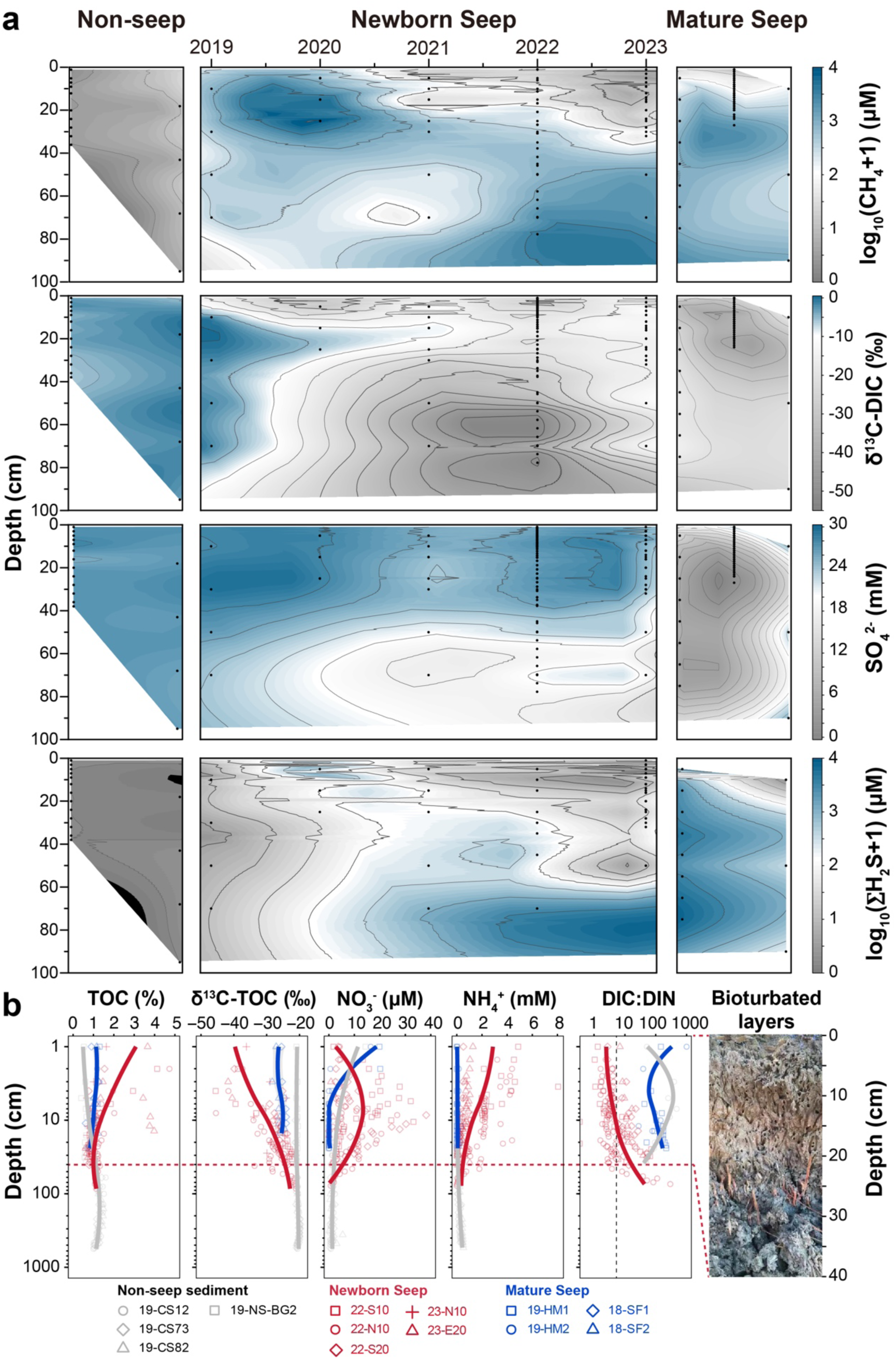
Time-series geochemical profiling reveals rapid methane removal and co-occurred active carbon, nitrogen, and sulfur cycling in the Newborn Seep sediments. **(a)** Contour plots of methane concentration (CH_4_), δ^13^C of dissolved inorganic carbon (δ^13^C-DIC), sulfate concentration (SO_4_^2-^), and total concentrations of hydrogen sulfide (∑H_2_S) in 0-100 cm sediments. Concentrations of CH_4_ and ∑H_2_S are log_10_(x+1) transformed, to better visualized the wide ranges of concentration gradients. Black dots indicate the data points. **(b)** Depth profiles of total organic carbon (TOC), δ^13^C-TOC, nitrate (NO_3_^-^), ammonium (NH_4_^+^), and ratios of dissolved inorganic carbon to nitrogen (DIC:DIN) at the Newborn Seep, measured in 2022 and 2023 when the macrofauna bioturbation activities maximized. The symbols and lines represent the measured data and smooth fitting using the ‘loess’ function, respectively. The photograph on the right shows the sediment profile of a box core with dense faunal burrows.

## Methane oxidations fueled by multiple electron acceptors

Targeted metagenomic and metatranscriptomic analyses performed during the period with intense bioturbation (2022–2023) confirm that methane oxidation was supported by a diverse suite of electron acceptors, underpinning the high metabolic activity and versatility of the Newborn Seep communities (Fig. 5). Metagenomic profiling combined with hierarchical clustering analysis reveals distinct metabolic traits across the non-seep, Newborn Seep, and mature seep ecosystems (Fig. 5a). Matching the geochemically inferred co-occurrence of diverse redox reactions in the strongly bioturbated layers (0-50 cm) of Newborn Seep, functional genes of aerobic methane/alkane oxidation and nitrate reduction were all relatively enriched and cooccurred with relatively low abundances of sulfate reduction and AOM genes. Noteworthily, metabolic traits of the deepest samples retrieved from the Newborn Seep were similar to those of the Mature Seep samples, suggesting a rapid functional succession of the leakage-impacted community toward mature seep community.

**Fig. 5.**
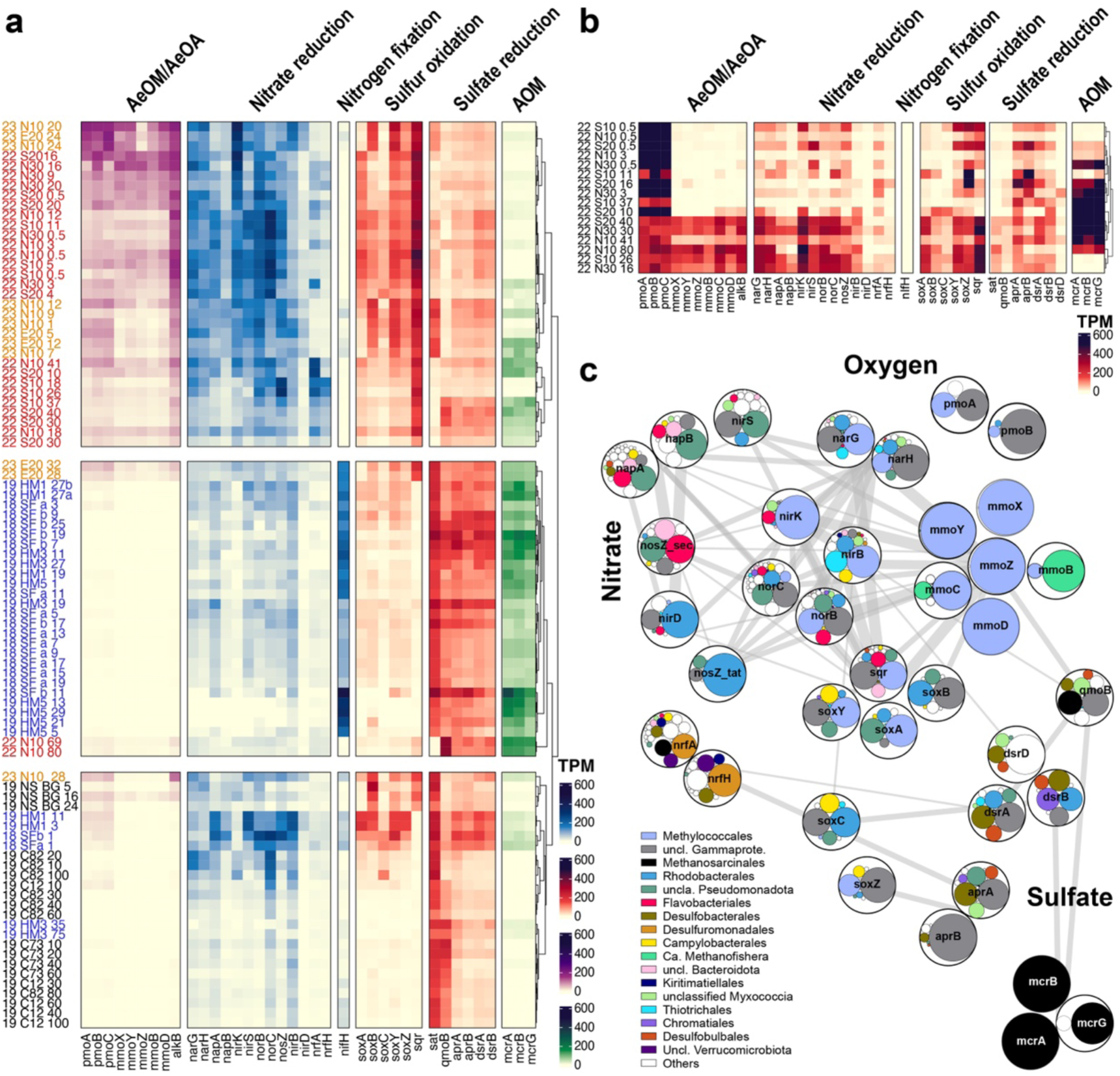
Metagenomic and metatranscriptomic analyses reveal active methane oxidations fueled by multiple electron acceptors at the Newborn Seep. Heatmaps showing the abundance of present (metagenomes, **a**) and transcribed (metatranscriptomes, **b**) key functional genes involved in aerobic methane/alkane oxidation (AeOM/AeOA), nitrate reduction, sulfur oxidation, sulfate reduction, and anaerobic methane oxidation (AOM). The color scale indicates transcripts per million (TPM), with darker colors reflecting higher abundance or transcription levels. Each row represents a sample, while each column corresponds to a specific gene or gene set. The names in red, orange, blue, and black colors indicate samples collected from the Newborn Seep in 2022, Newborn Seep in 2023, mature seeps, and non-seep background sediments, respectively. Nomenclature of sample names: sampling year_sampling site_sediment depth. (**c**) Gene co-transcription network linking major metabolic functions and microbial taxa. Each node (outer circle) represents a gene and each edge denotes significant correlations (Spearman *p*<0.05). Within each outer circle, the inner circles indicate the taxonomic composition associated with each gene. The network underscores the potential coupling of methane oxidations with the reductions of dioxygen, nitrate, and sulfate at the Newborn Seep.

Transcriptomic profiling and gene co-transcription network further indicate the potential coupling of methane oxidations with the reductions of dioxygen, nitrate, and sulfate in the Newborn Seep sediments (Fig. 5b, c). Transcript abundances of aerobic methane oxidation genes (*pmo*ABC) were high in all of the Newborn Seep samples examined, particularly in the 0-20 cm layers (Transcripts Per Million (TPM) >600, Fig. 5b). Notably, high transcript abundances of *pmo*ABC and the AOM genes *mcr*ABG cooccurred in 9 out of the 16 transcriptomes analyzed, even in the 0-20 cm sediments, matching the co-existence of aerobic and anaerobic methanotrophic clades in >85% of the Newborn Seep samples analyzed. This is most likely related to the dense faunal populations, which created spatiotemporal mosaics of oxic-anoxic microenvironments that facilitate the co-occurrence of various aerobic and anaerobic processes^46^. Intriguingly, another set of aerobic methane oxidation genes *mmo*XYZBD and nitrate reduction genes (*nar*GH, *nirBK*, *nor*BC, *nos*Z) were concurrently transcribed mainly in the 20-50 cm sediment layers (TPM 70-300) and thus strongly correlated (all with Spearman *r*>0.8, *p*<0.01; Fig. 5b, c). Transcripts of *mmo*XYZBD and partial denitrification genes (*nar*GH, *nirK*, *nor*BC) moreover, were all detected in *Methylococcales* (mainly *Methylomonadaceae*), indicating the potential coupling of these processes mediated by *Methylomonadaceae*. Collectively, it appears that the reductions of diverse electron acceptors (O_2_, NO_3_^-^, SO_4_^2-^) generally co-occurred and may have jointly fueled the highly active methane oxidations in surface sediments of the Newborn Seep.

## Implications

Our findings demonstrate that abrupt methane leakage triggers a rapid reorganization of seabed ecosystem, simultaneously causing a collapse of native biodiversity and the swift assembly of an effective “methane biofilter”. The majority of the benthic prokaryotic and eukaryotic organisms prove highly vulnerable to sudden methane influx. The established correlation between *in situ* methane concentration and biodiversity suggests that even micromolar-level methane increases can cause significant biodiversity loss, offering a potential benchmark for assessing the ecological risks in hydrate-rich seabed. Contrary to the paradigm of inherently slow deep-sea metabolisms^47,48^, we show that certain microbial and faunal groups can exploit sudden energy inputs with remarkable speed. Most notably, the anaerobic methanotroph ANME-3 responded even faster than its aerobic counterparts, a finding that challenges the established model of a multi-decadal delay in forming an anaerobic methane sink^14,24,49^. This suggests ANME-3 may act as a ‘pioneer species’ in emerging methane seeps, a trait warranting further physiological and metabolic investigations^50^. Continuous monitoring of this model ecosystem will reveal how and how fast the dominance of pioneering microbial and macrofaunal responders progressively shifts toward typical mature seep communities while maintaining chemosynthetic ecosystem functioning, thereby addressing key knowledge gaps regarding the formation and evolution of marine cold seeps.

The rapid formation of chemosynthetic ecosystem following methane leakage offers new perspectives on deep-sea carbon cycling. The newly-formed ecosystem demonstrates remarkable methane oxidation capacity, reaching rates among the highest recorded at natural seeps worldwide within just a few years^22,23^. Such exceptional methane removal capacity is largely sustained by the rapidly established animal-microbe synergies, in which opportunistic polychaetes induce redox fluctuations that facilitate the coupling of microbial methane oxidation with various redox reactions, substantially reshaping the carbon flows in deep-sea food webs (Fig. S2). In light of this animal-microbe interaction, the previously estimated 60-100 year “time window” for significant methane release prior to the establishment of an efficient methane biofilter^14,24,25^, may be significantly shorter. These new perspectives about the timescale and nature of seabed responses to methane leakage are imperative, particularly as thousands of offshore gas and oil wells worldwide show high rates (10-43%) of integrity failures that pose methane leakage risks^51,52^, and as subsurface methane reservoirs are increasingly destabilized under climate- and human-induced perturbations^6–9^. These insights are essential for refining the current models and predictions of how seabed ecosystems—and their vital role in mitigating methane and other hydrocarbon releases—will respond to escalating stressors from climate change and expanding industrial activities in the deep ocean^8,53,54^.

## Method

### 3D seismic survey

High-resolution 3D seismic survey was conducted by China National Offshore Oil Corporation in the Songnan Low uplift of Qiongdongnan Basin, South China Sea, to identify the seismic response characteristics associated with subseafloor geological structure, fluid migration, and gas hydrate accumulation. The data were acquired using 12 parallel streamers with 100 m spacings, with inline (NW-SE) and crossline (SW-NE) spacings of 12.5 m and 12.5 m, respectively, and 1.0 ms sampling interval. The data were processed as previously described^55^.

### Multibeam echosounder survey

Annual hydroacoustic surveys were conducted to track the activity of methane discharge from seafloor in the study region. Acoustic backscatter data from mid-water reflectors were collected using a shipborne Kongsberg EM302 multibeam echosounder, operating at a nominal frequency of 30 kHz with a swath width of 120°. The acoustic backscatter data were processed using CARIS 8.1 software to identify and extract gas flares.

### Sediment and porewater sampling

Sediment samples were collected from three major types of habitats in the South China Sea: the “Newborn Seep” (NBS)”, mature cold seeps (S18, ‘Haima’, and ‘Site F’), and non-seep continental slope sites (SCS12, SCS73, SCS82) (Fig. 1). Specifically, sediments were sampled from >40 sites around discharge center of the Newborn Seep during 2019–2023 (Fig. S1). Two non-seep background sites 19-NS-BG1 and 19-NS-BG2 were taken ∼200 m away from the discharge center in 2019, before the spreading of leakage on seafloor. Sediments were sampled primarily through push-core samplers of the Remotely Operated Vessel (ROV) “Haima”, compensated by piston core and box core sampling for subsurface geochemical profiling and macrofauna analysis, respectively. Porewater was extracted from sediment cores at different depths using the Rhizon sampler (0.15 μm pore size, Rhizosphere, Netherlands). Sediment cores were subsequently sliced into layers by every 2-5 cm. All samples were stored in proper temperatures and conditions for different analytical purposes.

### Macrofaunal analyses

Macrofaunal identification and quantification were performed using a combination of the sediment sieving-based method and image-based analysis. Sediments from box or push cores were sieved through 0.5-mm mesh, and the retrieved material preserved in 4% formaldehyde. Faunas were separated from debris in the laboratory and stored in 70% ethanol for later counting and identification. Notably, only few copepods can be retrieved by the sieving method despite their relatively high abundances recorded by underwater camera. The abundance of these faunal species therefore was estimated by analyses on images/videos that clearly visualized the individuals of copepods.

### Geochemical analyses

Methane concentrations were measured using a gas chromatograph (Shimadzu) equipped with a Barrier Discharge Ionization Detector, with detection limit of 0.01 μmol L^−1^. DIC concentration and its carbon isotopic ratio (δ^13^C_DIC_) were determined using Gas Bench II coupled with a Stable Isotope Ratio Mass Spectrometer (Delta 253plus, Thermo Scientific). Hydrogen sulfide was measured by methylene blue spectrophotometry (DR5000, Hach), while sulfate was quantified by ion chromatography (Dionex ICS-5000+, Thermo Scientific). NO_3_^-^, NO_2_^-^ and NH_4_^+^ were measured by using a continuous flow Auto Analyzer (AA3, SEAL), with detection limits of 0.015, 0.015 and 0.04 μmol L^−1^, respectively. TOC, TN, and the stable carbon isotope ratio of TOC (δ^13^C-TOC) were determined after removing the inorganic carbon through vaporization with concentrated HCl, using an elemental analyzer (Vario EL III) coupled to an isotope ratio mass spectrometer (Isoprime, Elementar) at the instrumental analysis center, Shanghai Jiao Tong University.

### Time-dependent reaction-transport model

To model the reaction rates and fluxes of dissolved chemical species in sediments, the evolution of porewater profiles over time is simulated using a 1D time-dependent reaction transport model:

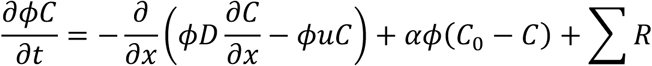

where *C* is the solute concentration (M L^-3^_fluid_), *t* is time, 𝜙 is porosity, *D* is the tortuosity-corrected diffusion coefficient^2^, *u* is the advection velocity, α is a nonlocal transport coefficient, *C_0_* is the concentration in the overlying water, and *ƩR* is the net rate of production or consumption of chemical *C* (M L^-3^_total_ T^-1^). The model takes into consideration dissolved oxygen, sulfate, methane, nitrate, ammonium, total sulfide, and dissolved inorganic carbon. Their concentrations are fixed at the upper boundary representing the overlying water (O_2_ = 250 µM, SO_4_^2-^ = 29 mM, NO_3_^-^ = 40 µM, NH_4_^+^ = 0 µM, TS = 0 µM, DIC = 2.2 mM, CH_4_ = 0 mM). The model accounts for reactions involved in the mineralization of organic matter (R_aer_, R_DNF_, R_DNRA_, R_SR_, R_MOG_; M L^3^_total_ T^-1^) and secondary reaction (R_no3ts_, R_sox_, R_nitri_, R_ch4o2_, R_aom_, R_no3ch4_; M L^-3^_fluid_ T^-1^). The rate parameters are primarily based on Wang and Van Cappellen (1996)^56^. The organic matter mineralization rate R_C_ is imposed as a function of depth, 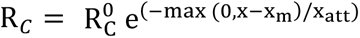, with 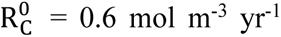. The irrigation coefficient was given the same functional depth dependence i.e., 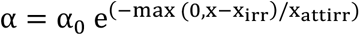, with α_0_= 200 yr^-1^ was set^57^.

Model simulations were carried out for a multi-year period capturing the evolution of the porewater profiles, using the R package “ReacTran” v1.4.3. First, steady state profiles were simulated during non-seep conditions, with 𝑢 = 𝑢_compaction_ = 𝑣_∞_ 𝜙_∞_⁄𝜙, where 𝑣_∞_ is the sedimentation rate solids at depth (set to 2 mm/yr). Mixing was set to be relatively shallow with x_irr_ = 0.03 m and x_attirr_ = 0.01 m, and mineralization rates decayed below the shallow mixing depth (x_m_ = 0.03 m; x_att_ = 0.01 m). These porewater profiles were then used as initial conditions for a 10-year simulation (Newborn Seep stage) with seepage, deeper solute mixing (x_irr_ = 0.3 m and x_attirr_ = 0.1 m) and mineralization extending deeper reflecting the effect of deep bioturbation (x_m_ = 0.3 m), to reflect the deeper mixed layer (Fig. 4). Subsequently, simulations were run toward steady state with continued seepage, but reduced mixing x_irr_ = 0.03 m and x_attirr_ = 0.01 m) and shallower mineralization (x_m_ = 0.03 m), representing the conditions found in nearby mature seeps. In the absence of seepage, a no concentration gradient condition was imposed at the lower domain boundary at a sediment depth of 2 m. During seepage, a flux was imposed as 𝐹 = 𝜙𝑢𝐶_6770_, where 𝑢 = 𝑢*_compaction_* _+_ *u_seepage_*, with *u_seepage_* = -0.25 m/yr. The seepage concentrations C_seep_ were set to O_2_ = 0 µM, SO_4_^2-^ = 0 mM, NO_3_^-^ = 0 µM, NH_4_^+^ = 20.2 µM, TS = 860 µM, DIC = 2.3 mM, CH_4_ = 50 mM. In addition, during seepage periods, it was assumed that a small amount of methane can partition from the rising gas phase into the porewater, adding to the dissolved methane concentration depending on the extent of undersaturation of the dissolved methane: *R_g_* = *k_g_*\*([CH_4_]_seep_ – [CH_4_]), with *k_g_* arbitrarily set to 0.044 yr^-1^.

### Monod biomass-explicit model

Methane oxidation rates were further estimated using the Monod biomass-explicit model, assuming the constant coupling between cell growth rate and specific substrate oxidation rate (i.e., there is no temporary uncoupling or ‘unproductive’ oxidation)^26^:

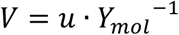

where 𝑉 is the specific rate of substrate oxidation (mol substrate oxidized per cell dry weight and time). 𝑢 is the net cell growth rate, estimated based on the temporal changes of 16S rRNA gene abundances of specific methanotrophic lineages (details see the method section “*In situ* cell growth”). 𝑌*_mol_* is the molar growth yield, was set to 0.6 g cell dry weight per mol CH_4_ oxidized according to the experiment data of typical methane seep sediments^26^. The derived specific rate of substrate oxidation 𝑉 was further converted to flux based on the mean abundances of methanotrophic lineages (10^7^–10^8^ cell g^-1^) and depth integration.

### Isotopic labeling experiments

Potential rates of denitrification were measured with ^15^N tracer slurry incubations. The fauna-free sediments were 1:7 diluted with artificial seawater, purged with helium for 30 min and then distributed into 12-mL Exetainers that were filled without headspace. The vials were preincubated at *in situ* temperature for 24 h and then spiked with ^15^NO_3_^-^ to a final concentration of ∼100 μM. Half vials were immediately treated with 200 μL 50% ZnCl_2_ solution to terminate the reaction and sampled as time zero, while the remaining vials were sampled after 24 hours. The rates of denitrification were calculated from ^29^N_2_ and ^30^N_2_ generation within the vials, which were measured by membrane inlet mass spectrometry^58,59^.

### Nucleic acids extraction and sequencing

Genomic DNA was extracted from 0.5-2.0 g sediments and DNA-free water (blank control) using a modified SDS-based extraction method^60^. For 16S rRNA gene sequencing, the hypervariable V4 region of 16S rRNA genes was amplified using the primer pair 515F (5’-GTGYCAGCMGCCGCGGTAA-3’) and 806R (5’GGACTACNVGGGTWTCTAAT-3’) and sequenced via the Illumina Miseq platform (Illumina, USA) as previously described^42,61^. For metagenomic sequencing, after library construction using the TruePrep DNA Library Prep Kit (Vazyme, China), paired-end sequencing with 150-bp length was performed at the Personalbio (Shanghai, China) using the Illumina NovaSeq platform. For metatranscriptomic sequencing, RNA of 17 selected samples was extracted from ∼5 g sediments using the RNeasy PowerSoil Total RNA Kit (QIAGEN, Germany). After removing ribosomal RNA and DNA by the Ribo-off rRNA depletion kit (Vazyme, China), libraries were prepared using the TruePrep RNA Library Prep Kit and sequenced on the Illumina NovaSeq-PE150 platform at the Personalbio.

### 16S rRNA gene sequence analysis

The raw reads of V4 region of 16S rRNA gene were processed and analyzed using the QIIME 2 platform v2020.11^62^. The primers and adapters were first trimmed out using Cutadapt v3.1^63^. Raw sequences were then processed using DADA2^64^, including quality filtering, denoising, paired-end sequence merging, chimera filtering and producing amplicon sequence variants (ASVs) and ASV Table. Taxonomy was assigned using q2-featureclassifier (a scikit-learn naive Bayes machine-learning classifier)^65^ with Silva database release 138^66^. Multiple sequence alignment and phylogenetic tree construction were performed using the QIIME 2 plugin q2phylogeny (align-to-tree-mafft-iqtree). Unassigned sequences, singletions and sequences affiliated with eukaryotes were discarded.

### Metagenome assembly and binning

Adapter and low-quality reads removal were performed by Trimmomatic v0.38^67^. Clean reads from each sample were *de novo* assembled using MEGAHIT v1.2.9 with a k-mer size of 4^68^. After removing contigs shorter than 2 kbp, binning was performed using MetaBAT2 v2.12.1^69^, MaxBin v2.2.7^70^, and CONCOCT v1.1.0^71^ with their default settings. Metagenome-assembled genomes (MAGs) from each assembly were dereplicated with DAS Tool v1.1.6^72^ and refined using RefineM v0.1.2^73^. Completeness and contamination of each MAG were evaluated using CheckM2^74^, retaining MAGs with >50% completeness and <10% contamination for further analysis. The taxonomy of each MAG was assigned by GTDB-TK v 2.1.1 with reference to GTDB R07-RS207^75^. The relative abundance of each MAG was calculated using CoverM with “genome” mode.

### Functional annotation

For >2 kbp contigs of each sample, gene calling was performed using Prodigal (-p meta) ^76^. For dominating MAGs (>1%), open reading frame was predicted from each MAG by Prodigal v2.6.3 with single mode^76^. Protein-coding genes and proteome of each MAG were annotated with the PFAM database using HMMER v3.3.2^77,78^, the KEGG database using Kofamscan v1.3.0^79,80^, and the eggnog database v5.0 using eggNOGmapper v2.1.12^81,82^. The resulting annotation for specific key genes was further verified across at least two databases or manually checked with custom database using Diamond blastp v0.9.14^83^. Genes encoding carbohydrate degradation enzymes described in the CAZymes database v10^84^ were searched using eCAMI with default 8 k-mer peptide library^85^, and the annotated CAZymes subfamilies were further mapped to the dbCAN2 database for substrate annotation^86^. The peptidase and proteinase encoding genes were annotated in the MEROPS database v12.4^87^ using Diamond blastp with a threshold of coverage >30% and e value < 1 × 10^−10^. The abundance of each gene in metagenomes was calculated as by counting the number of mapped reads for each gene using featureCount and normalized to Transcripts Per Million (TPM) value based on the gene length and sequencing depth^88^.

### Metatranscriptomic analysis

Metatranscriptomic reads were first quality filtered using Trimmomatic v 0.38^89^ and mRNA sequences were obtained by removing rRNA sequences with SortMeRNA v 4.3.6^90^ against both the SILVA^66^ and the default databases. Sam files were generated by mapping mRNA sequences to MAGs using hisat2 v 2.2.1^91^. Gene transcription was calculated by counting the number of unambiguously mapped reads for each gene using featureCount^88^. To compare transcription levels between genes, read counts were converted to TPM. A metabolic network at the transcriptional level was constructed based on the pairwise Spearman correlations between genes. Only correlations with *p*<0.01 were shown in the network. The taxonomic information of each gene was further annotated with the NR database using diamond blastp according to the best-hit target and visualized using the “circlify” package in Python (https://www.python.org).

### Quantitative PCR

Abundances of bacterial and archaeal 16S rRNA genes were quantified on a StepOne Plus (Applied Biosystems, USA) by SYBR-Green I-based quantitative PCR (qPCR). The primer pairs for Bacteria and Archaea were bac341f (5’-CCTACGGGWGGCWGCA-3’) / 519r (5’-TTACCGCGGCKGCTG-3’) and Uni519f (5′-CAGCMGCCGCGGTAA-3′) / Arc908R (5′-CCCGCCAATTCCTTTAAGTT-3′)^92,93^, respectively. The reaction mixture (20 μl) included 10 μl of ChamQ SYBR Color qPCR Master Mix (Vazyme, China), 1.6 μl (Bacteria) / 1.6 μl (Archaea) of each primer (10 μM), and 1 μl of template DNA. The thermal cycling program was: initial denaturation at 95 °C for 15 min, 40 cycles at 95 °C for 30 s, 56 °C (Bacteria) / 60 °C (Archaea) for 30 s, and 72 °C for 30 s. Plasmids of 16S rRNA genes from *Marinobacter* sp. and *Bathyarchaeota* were used as bacterial and archaeal standards. Microbial cell abundance was subsequently estimated using the average 16S gene copies per cell (Bacteria: 4.12 ± 2.75 copies cell^−1^; Archaea: 1.61 ± 0.88 copies cell^−1^) based on the rrn database (https://rrndb.umms.med.umich.edu/).

### *In situ* cell growth

The net cell doubling time (DT) was estimated as the quotient of time and log_2_-fold changes in 16S rRNA gene copies of specific lineages:

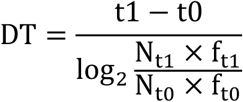

where t_1_ is the specific time point after the onset of methane seepage at t_0_. N is the total 16S rRNA gene copy number based on qPCR, f is the fraction (i.e., relative abundance) of specific lineage based on the 16S rRNA gene sequencing data. For N and f at t_0_, data retrieved from the non-seep background sites were used to represent the community status before methane leakage. The net cell growth rates (R) were further calculated as:

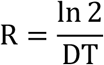

### Statistics

All statistical analyses were performed in R (https://www.Rproject.org). Microbial alphadiversity indices and PCoA coordinates using Bray-Curtis distances were calculated using the “phyloseq” package^94^. Using the “vegan” package^95^, we performed: 1) Anosim analysis to examine the community (dis)similarity between different groups of samples, 2) PERMANOVA analysis to examine the powers of different categorical and geochemical factors in explaining community variations, 3) two-sided Welch’s t test with Bonferroni-Holm correction to compared means between two independent groups without assuming equal population variances. The succession rates of microbial communities were calculated as previously described, by calculating the temporal variations of the community distances measured by both Bray-Curtis and Sorensen metrics with time^96^. The potential contribution of methane derived carbon to the faunal diet was calculated by the “MixSIAR” pakage, using the default uninformative prior and Markov chain Monte Carlo settings^97^.

## Data Availability

The data are available in the article or the online supplementary material.

## Acknowledgement

We thank the scientists and crew of the Haiyangdizhi VI and Haima ROV for collecting and processing sample and data; Dr. Fan Yang, Dr. Mingyang Niu, Lin Yuan, Ben Zhu, Jiahui Ji, Yifan Chen, Xiang Ji, Sihan Li from the Shanghai Jiao Tong University for help with the geochemical analyses. This work was supported by the following grants: the National Natural Science Foundation of China (Grant 42230401, 42276087, 42306057, and 42576121), Science and Technology Commission of Shanghai Municipality (Grant 23R1428500), Guangdong Basic and Applied Basic Research Foundation (Grant 2019B030302004), Marine Geological Survey Program of China Geological Survey (Grant DD20230065), Simons Foundation (Grant 824763 to S.E.R.), U.S. National Science Foundation (Grant 2220309 to C.M.), and U.S. DOE (Grant DE-SC0022991 to C.M.).

