## Supplementary materials for "Rapid ecosystem collapse and biofilter formation following seabed methane leakage"

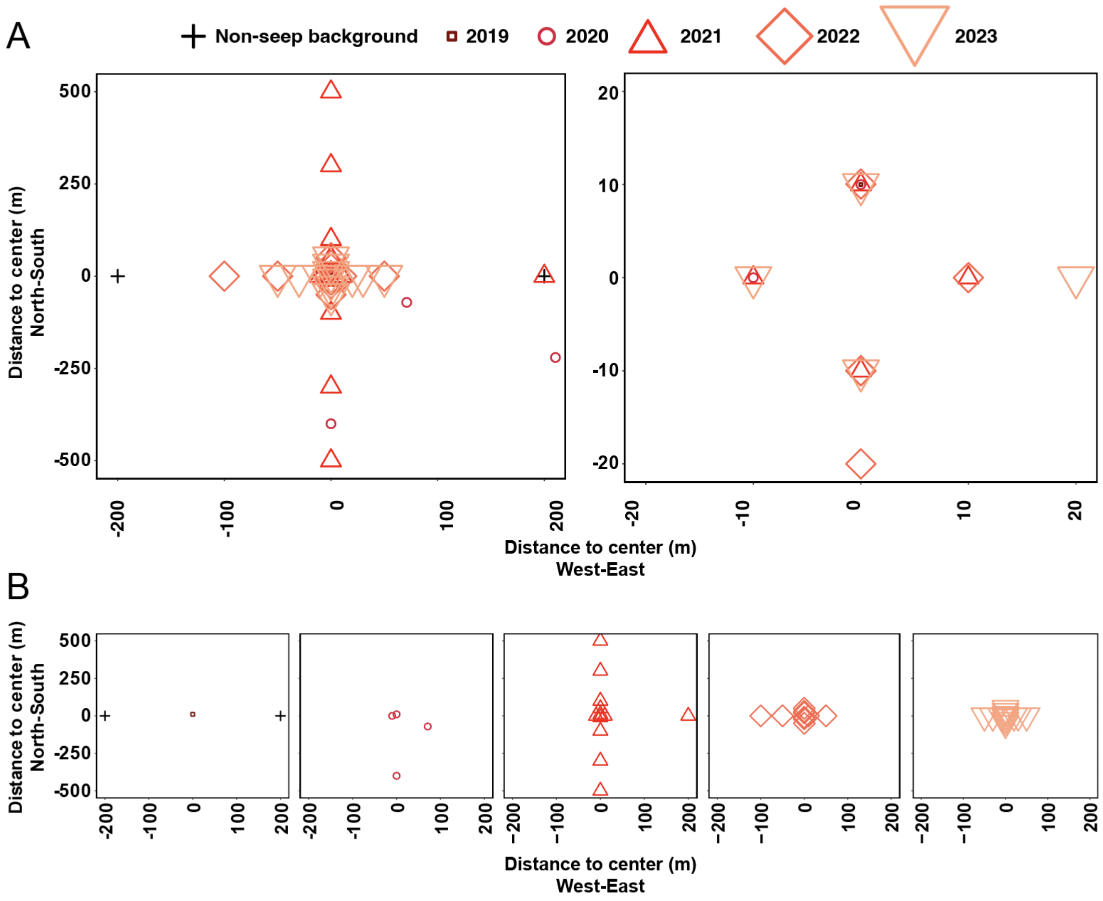


**Fig. S1.** Information of the sampling sites around the Newborn Seep, with coordinate (0, 0) denotes the discharge center. **(A)** All sampling sites viewed at different spatial scales. **(B)** Yearly sampling sites from 2019-2023. The non-seep background sites 19-NS-BG1 and 19-NS-BG2 were taken ~200 m away from the discharge center of Newborn Seep in 2019, before the spreading of leakage on seafloor.

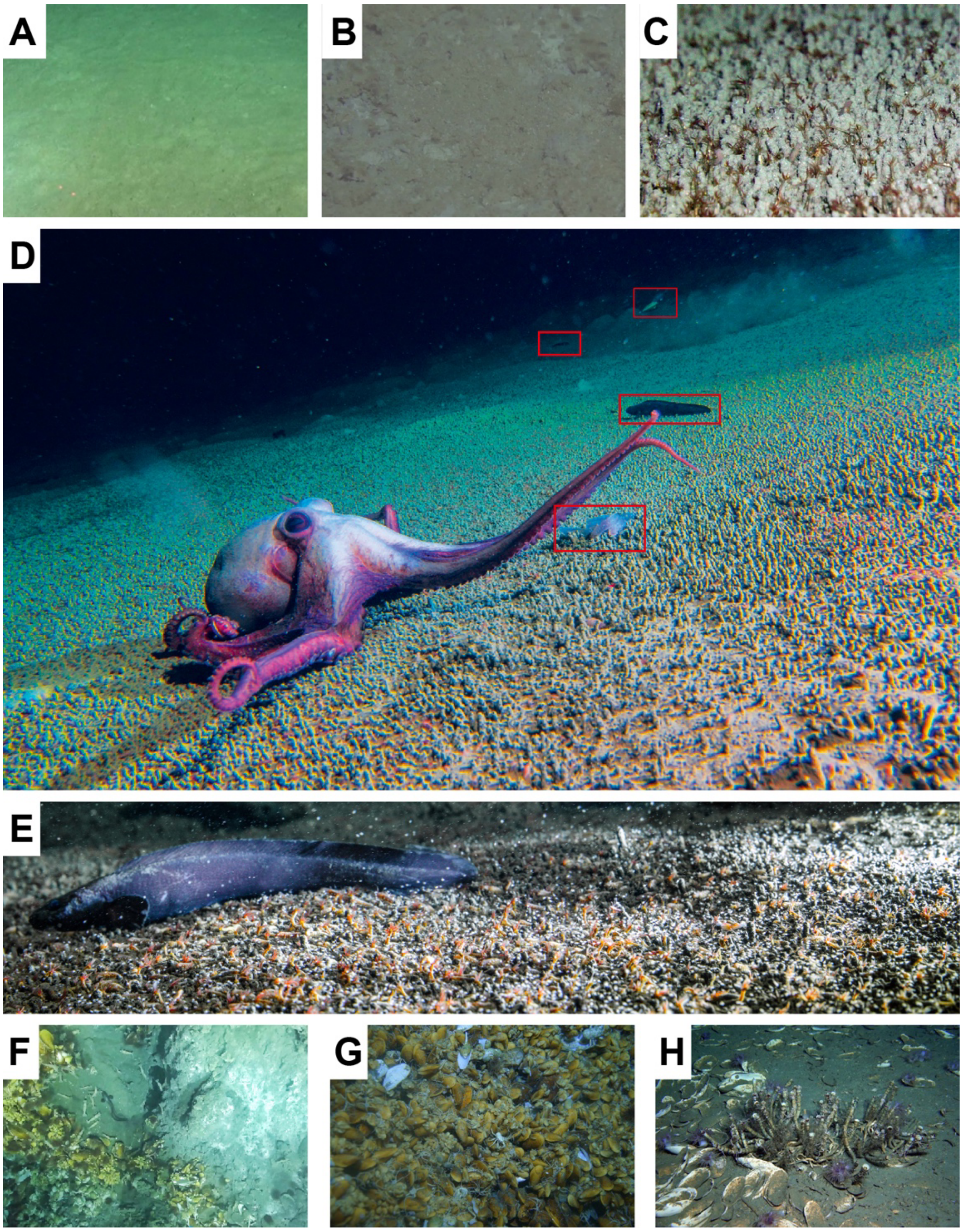


**Fig. S2.** In-situ ROV images showing active methane venting and associated seep fauna at multiple sites in the Qiongdongnan Basin (QDN) and Haima region. **(A-B)** the nearly 'bare' seafloor observed before (2018) and after the methane leakage in 2019, respectively; **(C)** a dense assemblage of polychaetes blanketing the soft sediments of the Newborn Seep observed in 2021; **(D-E)** higher trophic animals attracted by the newly formed biomass-rich ecosystem; **(F)** carbonate outcrops, white microbial mats, and mussel aggregations indicative of active seepage at the mature seep site S18; **(G-H)** typical mature seep fauna including the mussels, tubeworms, and clams at the mature "Haima" cold seeps.

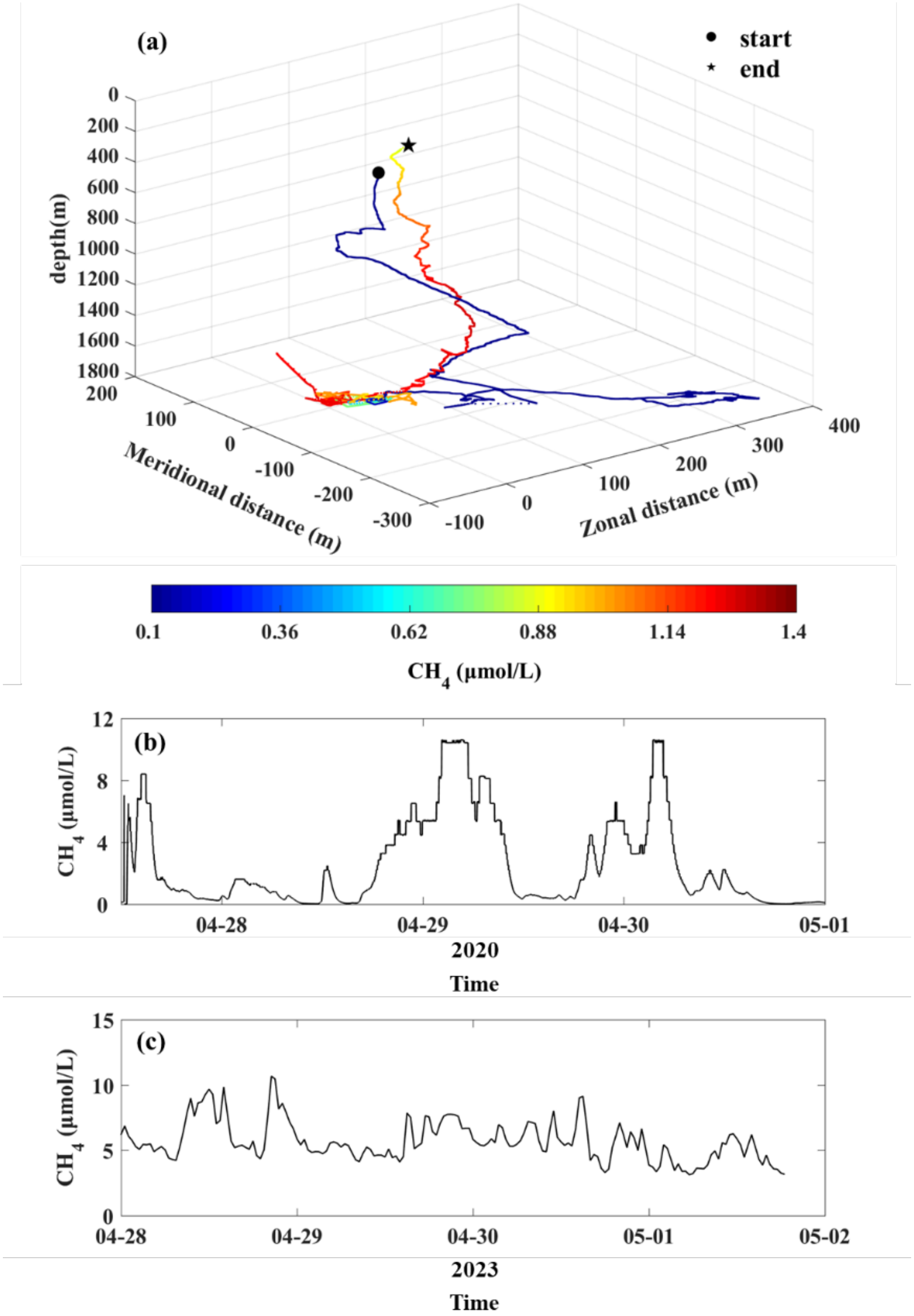


**Figure S3.** Temporal variation of seawater methane concentrations at the Newborn Seep site: **(a)** Three-dimensional plot of the sampling path, with meridional distance (m), zonal distance (m), and depth (m) on the axes. The color scale indicates measured $CH_4$ concentrations (μmol/L). The black symbols (circle, star) denote the start and end points of the survey. **(b)** Time series of methane concentrations ($CH_4$, μmol/L) recorded in water column above the discharge center, from April 27 to May 1, 2020. **(c)** Time series of methane concentrations ($CH_4$, μmol/L) recorded in water column above the discharge center, during a separate campaign in 2023 (April 28 to May 2).

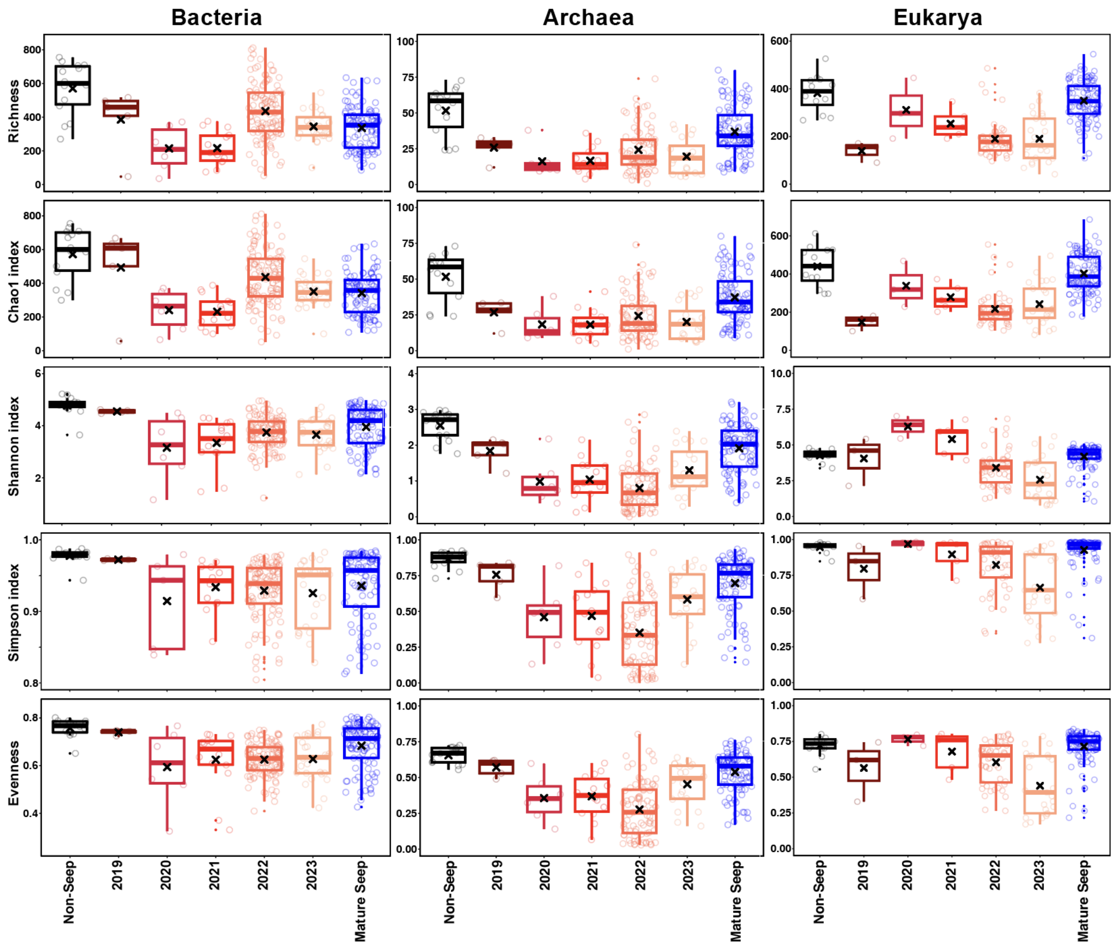


**Figure S4. Multi-year variations of the bacterial, archaeal, and eukaryotic diversity indices following methane leakage.** The 'non-seep seafloor' group include data from the adjacent non-seep background sites, while the 'mature seep' group include data from the 'Haima' and 'Site F' cold seeps in the South China Sea (see Supplementary Table 1).

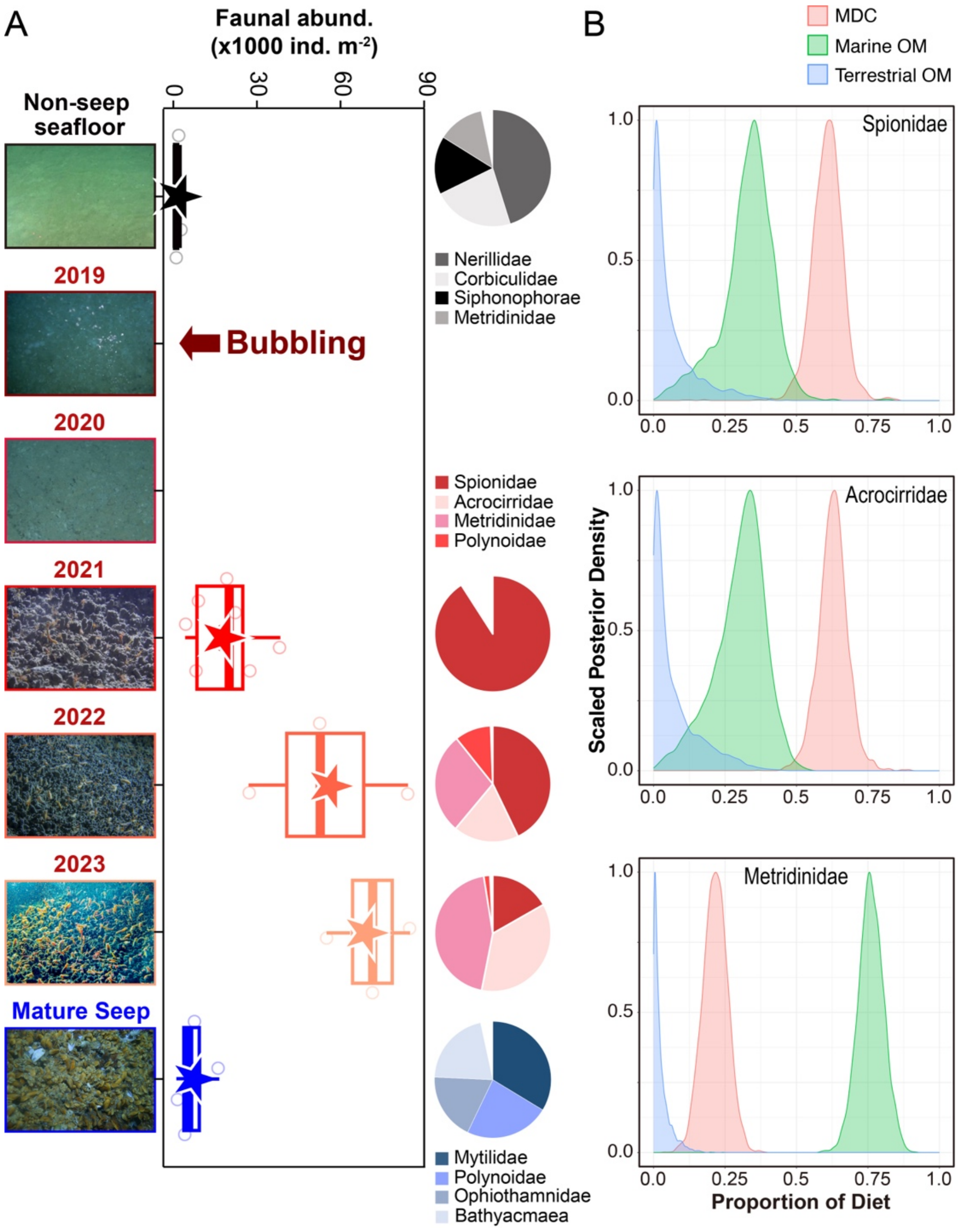


**Fig. S5. Rapid proliferation of faunal populations after methane leakage. (A)** Time-series changes in benthic faunal abundance ($\times10^3$ individuals $m^{-2}$) from a non-seep seafloor reference (top) through multiple post-discharge years (2019-2023) to a mature seep environment (bottom). The boxplots display medians, quartiles, and data ranges. The dots within boxplots denote the mean values. The pie charts illustrate the relative abundances of major faunal groups. Active bubbling from the seafloor was first observed in 2019, in line with the onset of methane discharge detected by multibeam echosounder. **(B)** Bayesian stable isotope mixing model results showing the proportional contributions of three dietary carbon sources—methane-derived carbon (MDC), marine organic matter (Marine OM), and terrestrial organic matter

(Terrestrial OM)—to major fauna groups (Spionidae, Acrocirridae, Metridinidae) residing at the Newborn Seep. The x-axis and y-axis denote the proportional contribution of each carbon source to the faunal diet and the scaled posterior density, respectively.

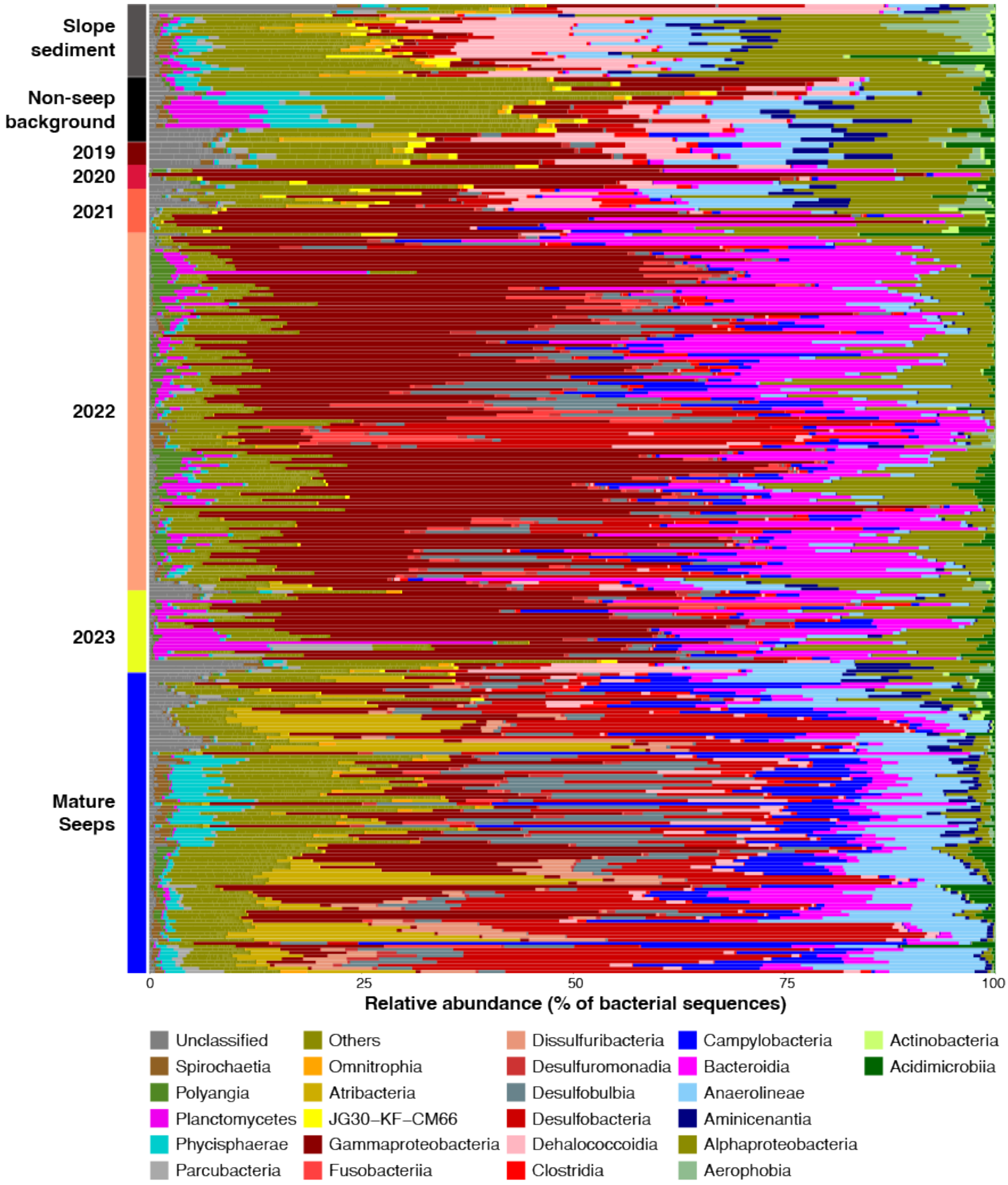


**Fig. S6.** Class-level bacterial community compositions in sediments across the continental slope, non-seep background, Newborn Seep (2019-2023), and mature seep sites.

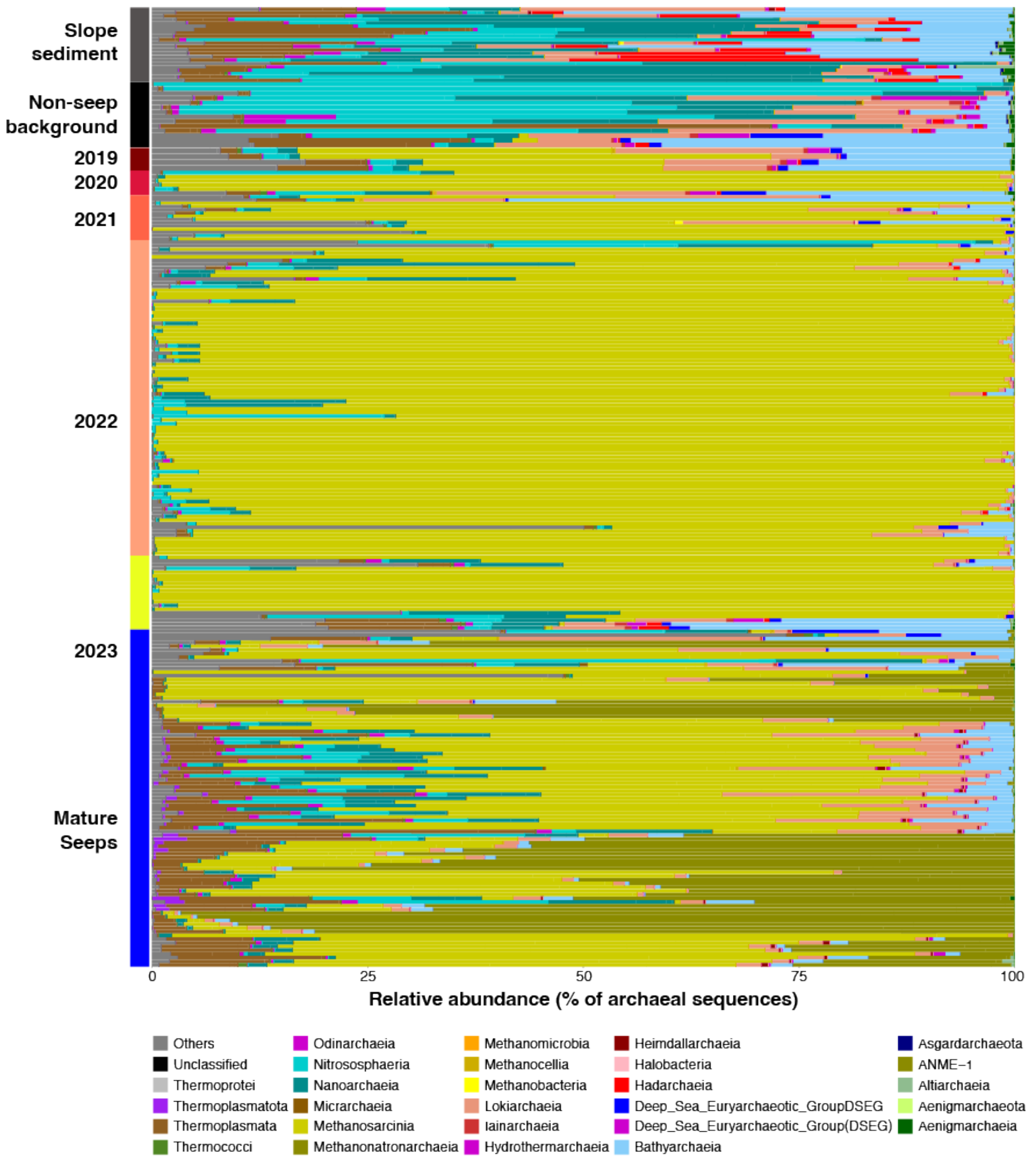


**Fig. S7.** Class-level archaeal community compositions in sediments across the continental slope, non-seep background, Newborn Seep (2019-2023), and mature seep sites.

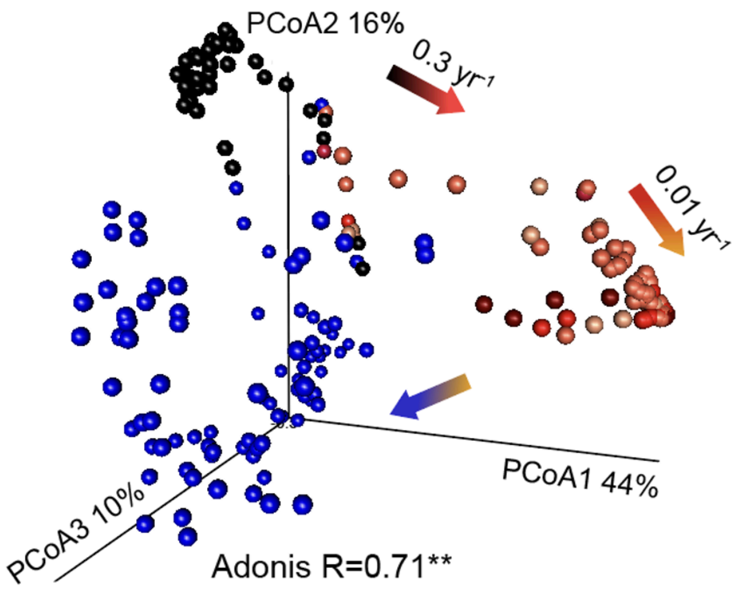


**Fig. S8.** Principal coordinates analysis (PCoA) plots based on archaeal 16S rRNA genes, demonstrating distinct archaeal community clustering by seep stage and year. The Adonis R and p-values indicate the proportion of variation explained by seep stage (**: $p$<0.01).

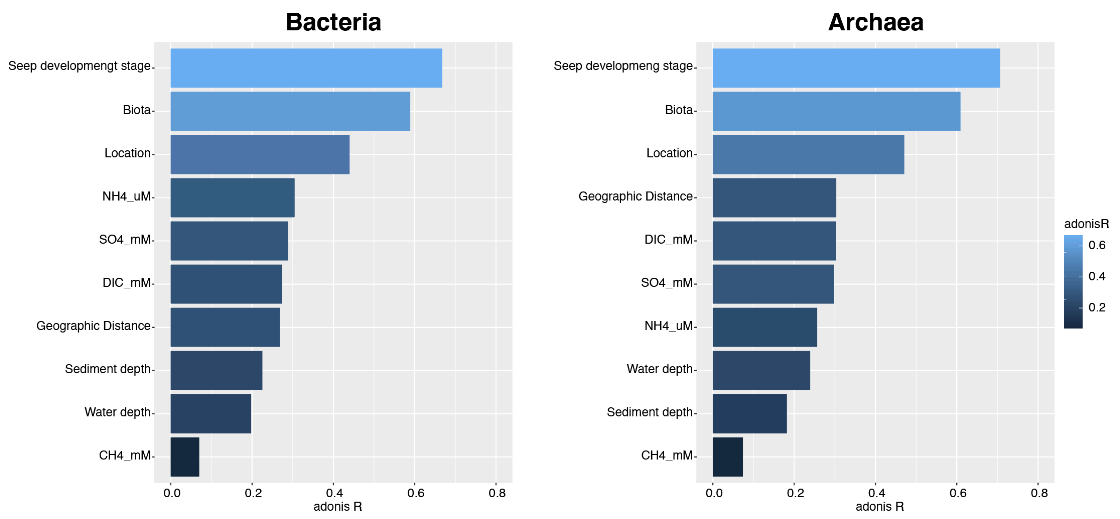


**Fig. S9.** Results of PERMANOVA (adonis) analyses showing the explanatory power (adonis R) of various environmental and spatial factors on sediment microbial community structure. Bars are sorted in descending order of their adonis R values, with "Seep development stage" exerting the strongest influence. Other factors include "Biota-," sampling "Location," ammonium (µM), sulfate (mM), methane (mM), dissolved inorganic carbon (mM), geographic distance, water depth, and sediment depth. The color intensity corresponding to the magnitude of the adonis R value.

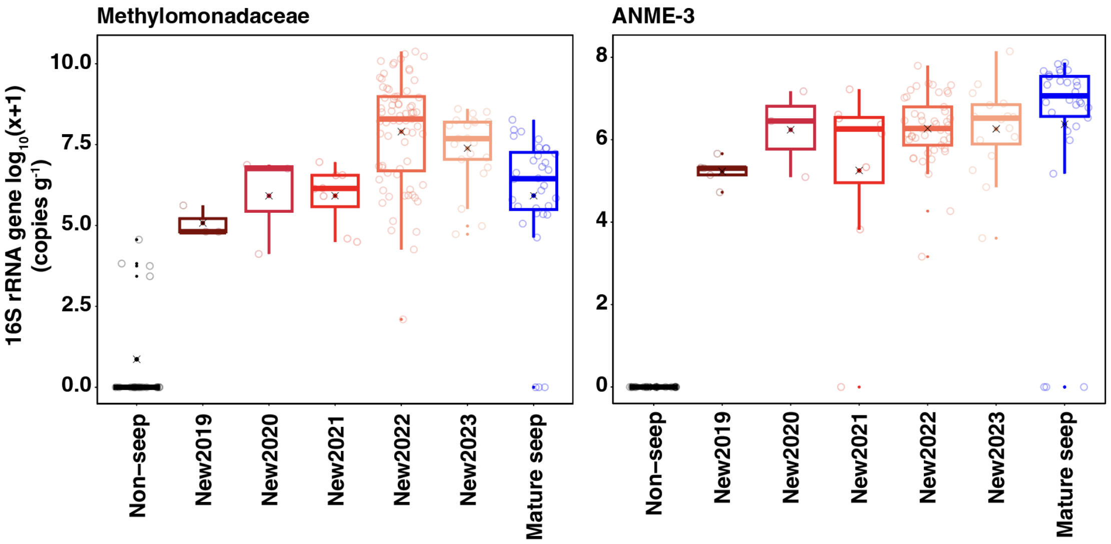


**Fig. S10.** Changes of absolute 16S rRNA gene abundances of representative aerobic (*Methylomonadaceae*) and anaerobic methanotrophic lineages (ANME-3) across the non-seep, Newborn Seep, and mature seep stages.
